# LoopSage: An Energy-Based Monte Carlo approach for the Loop Extrusion Modelling of Chromatin

**DOI:** 10.1101/2024.01.10.574968

**Authors:** Sevastianos Korsak, Dariusz Plewczynski

## Abstract

The connection between the patterns observed in 3C-type experiments and the modeling of polymers remains unresolved. This paper presents a simulation pipeline that generates thermodynamic ensembles of 3D structures for topologically associated domain (TAD) regions by loop extrusion model (LEM). The simulations consist of two main components: a stochastic simulation phase, employing a Monte Carlo approach to simulate the binding positions of cohesins, and a dynamical simulation phase, utilizing these cohesins’ positions to create 3D structures. In this approach, the system’s total energy is the combined result of the Monte Carlo energy and the molecular simulation energy, which are iteratively updated. The structural maintenance of chromosomes (SMC) protein complexes are represented as loop extruders, while the CCCTC-binding factor (CTCF) locations on DNA sequence are modeled as energy minima on the Monte Carlo energy landscape. Finally, the spatial distances between DNA segments from ChIA-PET experiments are compared with the computer simulations, and we observe significant Pearson correlations between predictions and the real data. LoopSage model offers a fresh perspective on chromatin loop dynamics, allowing us to observe phase transition between sparse and condensed states in chromatin.

## 1 Introduction

Chromatin exhibits a hierarchical organization, with various levels of structure governed by distinct biophysical principles [1–3]. At the fundamental micro-scale level, the basic DNA packing units, i.e. nucleosomes provide beads on a string microscopic nanometer-scale structure of chromatin [4–6]. Moving up on the organizational scale, chromatin fiber forms loops, flares (stripes) and topologically associating domains (TADs), otherwise named CCDs, chromatin contact domains) [7], which can be identified by the advanced experimental 3C-type techniques such as Hi-C, ChIA-PET, and Hi-ChIP. The formation of these meso-scale structures of various sizes are orchestrated by key protein factors, including cohesin or condensin SMC complexes and CTCF proteins [8]. Chromatin organization at macroscale involves the formation of A and B compartments, and their subcompartments [9]. The precise biophysical mechanisms underlying compartments formation remains elusive, although the prevailing theory suggests that epigenetic marks and histone modifications establish distinct nucleosomes populations with diverse physico-chemical properties [10]. Furthermore, chromatin is organized into distinct chromosomal territories, compacted all together within an approximately spherical container by nuclear membrane.

In this study, we concentrate on the spatial and temporal meso-scale of chromatin loops and TADs, where loop extrusion (LE) emerges as the dominant molecular mechanism of chromatin organization. According to the LE model, a loop extrusion factor (LEF), often an SMC complex, binds to chromatin and initiates the extrusion of a loop connecting different chromatin segments in sequential order, the next one after the previous one following the time course [11]. The boundaries of loop extrusion are determined by CTCF proteins called alternatively boundary elements (BEs), which binds to DNA at the specific sites encoded by the sequence motifs with two possible orientations [12]. The orientation of CTCF plays a critical role in the loop extrusion mechanism, as CTCF with incorrect orientation fails to impede the movement of LEFs.

Understanding the structural patterns observed in 3C-type experiments, which visualize the allagainst-all spatial proximities between genomic segments as heatmaps, represents a major challenge in 3D genomics. First, the existence and the characterization of the higher order organization of chromatin, and secondly deciphering its biological function in mammalian genomes [7]. The careful analysis of HiC heatmaps, which are *N* × *N* matrices of real numbers representing probabilities of contact between (*i, j*) pairs of genomic locations at the given resolution (eg. five thousands base pairs, 5kbp), reveals non-random distributions of structural patterns such as stripes (flares), TADs (chromatin contact domains), and chromatin loops, located differently relative to the diagonal (*i* == *j*) [8]. In the heatmap matrix *N* represent the number of 5kbp DNA segments, in other words bins, in the whole genome. Each of these observed spatial patterns serve its own functional purpose described by chromatin biology. However, establishing a causal connection between specific heatmap features and their underlying biophysical rationale remains elusive. The available experimental 3C-type data is often aggregated across population of tens or hundreds of millions of cells, and averaged over a cell-cycle. Chromatin loops are typically interpreted as regions, where LEFs remain associated to DNA for extended periods of time, stripes are associated with one-sided loop extrusion by cohesin binded to CTCF as a barrier, and TADs are linked to chromatin regions exhibiting high LEF activity between convergent CTCF sites. Moreover, CTCF proteins are enriched at TAD boundaries supporting their insulator function by blocking the communication between enhancers and promoters across the boundary, while promoting interactions within the chromatin contact domain.

Current state-of-the-art models of loop extrusion employ stochastic Monte Carlo methods combined with molecular simulations [13–19]. Two primary modeling approaches have emerged. First, some models investigate how loop extrusion impacts chromatin folding and the global chromatin structure, examining the effects of different types of LE (one-sided, two-sided, symmetrical, asymmetrical) [16, 18, 20] and exploring the influence of other protein remodelers, such as RNApII, on cohesin dynamics [21–24]. Second, other models aim to replicate experimental heatmaps obtained from 3C data by employing simplified biophysical assumptions regarding LEFs and BEs dynamics [13, 15, 17]. In these models, cohesins are often represented as particles with two degrees of freedom, undergoing one-dimensional diffusive motion along the linear DNA fiber to establish connections between distal chromatin regions. Convergent CTCF orientations are typically modeled as strong barriers, assuming that LEFs cannot pass through each other-an observation supported by experimental evidence [11]. Another crucial assumption concerns the reloading rate of each LEF, which governs its unloading from one chromatin fiber position and subsequent random loading at another genomic loci. Within the second framework, each LEF as a ring-like structure either topologically enforce the proximity of two chromatin fibers [25, 26], or forms covalent chemical bonds between the two chromatin points it connects [27, 28]. The long range contacts formed this way are believed to be sufficient to explain the TAD-scale chromatin structure observed in 3C-type experiments.

Despite the extensive use of stochastic simulations in chromatin research, several challenges are still present in the modeling of loop extrusion. One of the major challenges is the development of an energy-based stochastic model, akin to the Ising model [29, 30], capable of accurately capturing the motion of LEFs along the chromatin fiber using methods such as Metropolis or Simulated Annealing, which is equivalent to the solution of the inverse problem in statistical mechanics with unknown Hamiltonian [31]. However, the non-equilibrium nature of the models hinders the fulfillment of the condition of detailed balance [15]. Specifically, the probability in the master equation for a LEF to move towards a BE is considerably higher than the probability of moving away from it. Overcoming this issue requires careful consideration. One possible solution to address this challenge is to make specific assumptions regarding the motion of each LEF [13, 15, 19], as discussed earlier, in order to bypass the need for constructing a Hamiltonian. However, this approach lacks mathematical rigor and elegance. Additionally, determining the experimental parameters necessary for these simulations remains difficult due to limited empirical knowledge.

In this study, we propose a novel computational model that satisfies the condition of detailed balance by incorporating the CTCF binding locations as barriers directly into the system’s Hamiltonian, instead of including them in the master equation [15]. By employing this mathematical technique, it becomes possible to define Metropolis simulations capable of reconstructing realistic three-dimensional chromatin structures, that they represent a thermodymic ensemble of possible polymer conformations that are linked to a region of interest. Our approach utilizes a simplified framework that draws samples from a Boltzmann distribution governed by a Hamiltonian energy functional consisting of three terms: a folding or entropic cost, a crossing penalty, and a binding one. The main advantage of our model is its improved clarity in the implementation of LEM physics, allowing for a more precise specification of its biophysical parameters. This can guide us to the discovery of phase transitions and provide deeper insights into the fundamental principles underlying chromatin folding, in addition to previous research [32–36]. We hope that we are introducing a more mathematically consistent approach, with a shorter and more elegant programming implementation, which can help scientific community to have a better understanding of loop extrusion process.

## 2 Method

The methodology employed in this study consists of two distinct components: stochastic simulation of LEFs’ trajectories, and molecular dynamics 3D conformation generation. The stochastic simulation phase assumes that the positions of LEFs can be described by two degrees of freedom, namely *m*_*i*_ and *n*_*i*_, with the constraint that *m*_*i*_ *< n*_*i*_. These variables determine two anchors to which single LEF is connected. The primary objective of these simulations is to explore the phase-space of LEF positions as a function of pseudo-time, denoted as *m*_*i*_(*t*) and *n*_*i*_(*t*), and subsequently reconstruct realistic trajectories based on fundamental principles governing LEF kinetics.

Following the stochastic simulations, the obtained LEF positions are utilized as inputs for a molecular dynamics model, enabling the creation of three-dimensional structures. The resulting thermodynamic ensemble of these structures provides a realistic representation of our system. Notably, the average all-versus-all heatmap representing inverse of spatial distance derived from this ensemble exhibits similar characteristics to the experimental data, confirming its ability to capture relevant properties of the system under investigation.

### 2.1 Data processing and the genome-wide CTCF motif identification

For the purpose of this research, we utilized publicly available CTCF ChIA-PET data [37] of GM12878 cell line obtained from the ENCODE project [38] (check Data Availability section). Our focus was primarily on investigating chromatin loops within this dataset as they were extracted by ChIA-PIPE pipeline [39]. By chromatin loops we imply regions of chromatin fiber that they are in close topological proximity. Thus ChIA-PET loops provide us the information about the interacting regions, which are called anchors and an interaction strength called PET-count. Given the large number of ChIA-PET chromatin loops available (1181427 loops for the whole genome), we applied two filtering steps to narrow down our analysis. Firstly, we excluded loops with a PET-count of less than or equal to 10 focusing only on the strongest chromatin interactions mediated by CTCF protein. Subsequently, we applied a second filtering step to retain interactions that specifically represented CTCF loops, ensuring the presence of a CTCF motif at the respective anchors of each loop. The reason why we do that is because weak loops would constribute in our energy landscape mainly as noise, and we would like to assist our stochastic model to reach faster the local minima.

To identify CTCF motifs in DNA sequence, their positions and orientations, we employed an approach that considered CTCF ChIP-seq within the given genomic interval. To process the wholegenome sequence data as personalized genomes, we substituted the nucleotides from the reference genome with single nucleotide variants (SNVs) obtained from the VCF file of a given individual (in this case NA12878 from 1000 genomes project). This ensures us that the processed genome DNA sequence closely matched the provided individual sample rather than the reference one. Next, we identified motifs within the two anchor regions of each loop using these modified, i.e. personalized genome sequence, and the Bio.motifs package from Biopython [40]. Each motif occurrence was represented by its position and a score, measured with the logarithmic scale (log_2_ *n*), indicating the DNA sequence likelihood to bind CTCF protein compared to background noise. For instance, a score of 2.0 signifies that the identified motif occurrence is four times more probable to be the motif rather than background noise. We collected all motif occurrences within the anchors, considering only those with a minimum score of 2.0. The strengths of these motifs were calculated separately for forward and backward directions. From this data, we computed the relative strength of motifs in the backward direction by dividing the sum of backward motif strengths by the total sum of strengths. Similarly, by using 1 − *p*, we obtained the relative strength of motifs in the forward direction, where *p* represents the relative strength of backward motifs.

### 2.2 Stochastic Simulation (LoopSage)

The following three Monte Carlo (MC) moves are permitted for a specific LEF in our stochastic simulation model (Fig.1A),

1. Rightward sliding (*n*_*i*_ → *n*_*i*_ + 1 to the right).
2. Leftward sliding (*m*_*i*_ → *m*_*i*_ − 1 to the left).
3. Rebinding at a different location with initial loop length *< ℓ > /*8, where *< ℓ >* is the average loop length as it is estimated from the loop length distribution from the bedpe file representing ChIA-PET experimentally identified loops.

**Figure 1:**
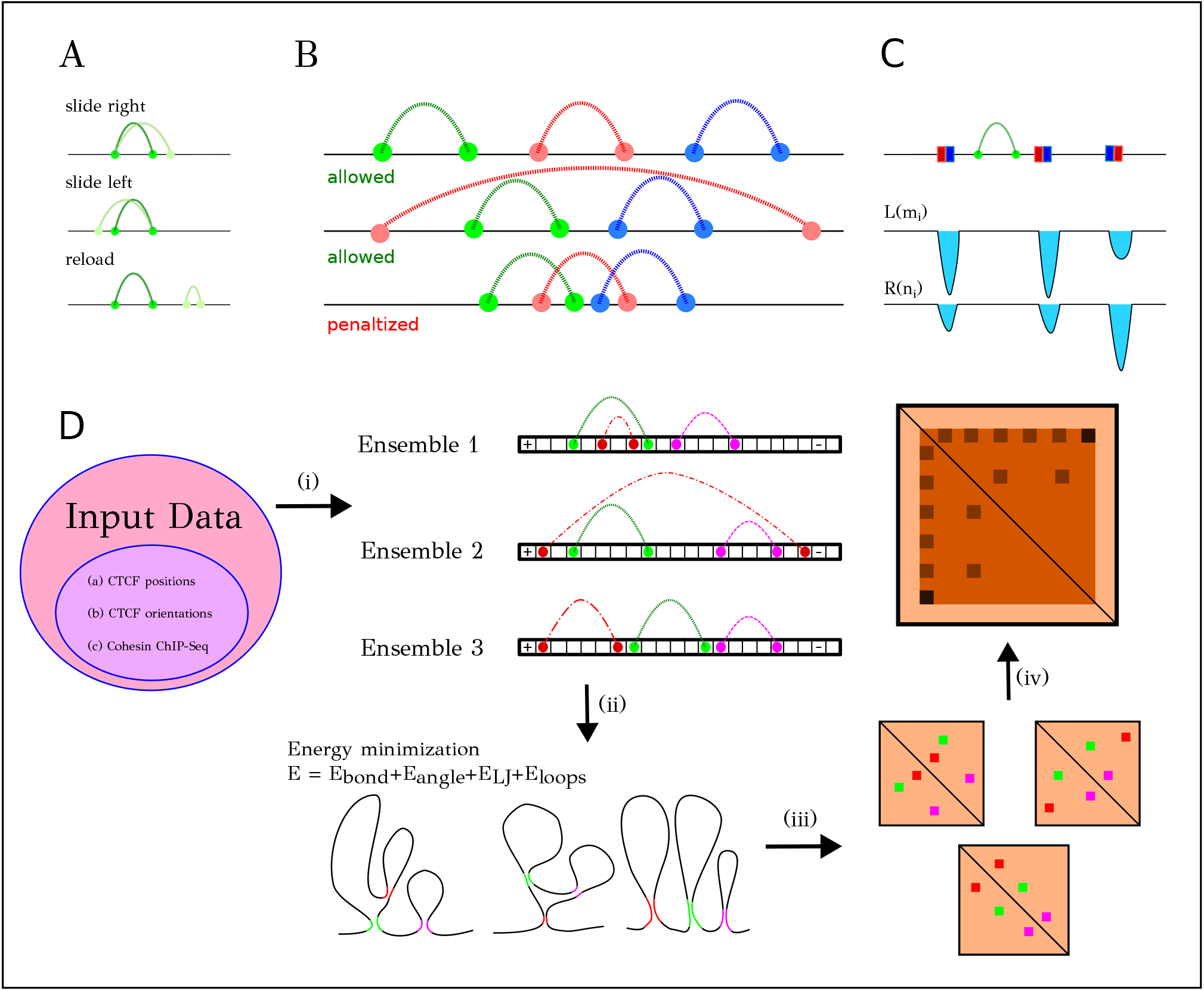
Description of the simulation method. (A) Possible Monte Carlo moves of the stochastic simulation. (B) Different conformation of loops. The first two ones are parallel and nested loops respectively and they are allowed. In the last case, there is conformation which is not allowed because it is responsible for crossing of chromatin. (C) CTCF points are linked to the energy minima of the two functionals *L*(*m*_*i*_), *R*(*n*_*i*_). (D) Stochastic simulation pipeline: (i) the simulation starts by processing the experimental data for CTCF positions and the orientation of the underlying binding motif on DNA sequence. SMC coverage file, from cohesin ChIP-Seq experiments, is an optional input for the model. Stochastic simulations generates an ensemble of LEF positions as output for further steps of the algorithm. (ii) After energy minimization the 3D structures that correspond to these LEF positions are computed. (iii) From these 3D structures, it is possible to compute the 2D inverse distance heatmaps of each one of 3D conformations. (iv) After averaging over many LEF occupancies and 3D models, we obtain an average inverse distance heatmap that should reproduce the experimental one.

Above MC movements are biophysically realistic and represent the basic linear relocation of LEFs along DNA chain, and three-dimensional redistribution of it. However, they do not satisfy any other constraints like the barrier effect of CTCF or other LEFs [11]. Therefore, we propose the energy function that permits biologically plausible moves. The energy function is defined as follows:

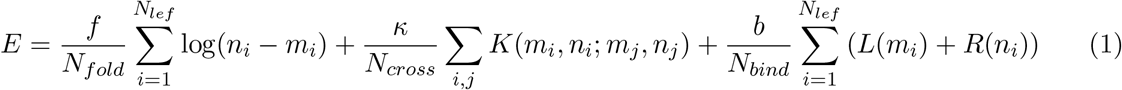

The first term represents the entropic cost [41] associated with chromatin folding, while the second term penalizes the occurrence of cohesin crossing (Fig. 1B). We define the function *K*(*m*_*i*_, *n*_*i*_; *m*_*j*_, *n*_*j*_) as follows:

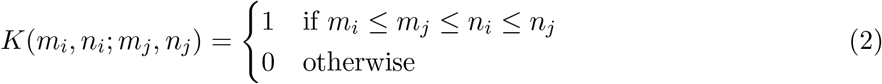

The functions *L*(·) and *R*(·) represent the binding potentials and are orientation-specific, distinguishing the left and right positions of LEF due to the orientation specificity of underlying CTCF binding motif on DNA sequence. Consequently, the presence of a gap in these functions indicates the presence of a CTCF site (Fig. 1C). These functions are derived from CTCF binning data, like CTCF loops from ChIA-PET data, and by running the script for probabilistic orientation. It is worth mentioning that the coefficient *f* should be chosen on the same order of magnitude as the total length of polymer, *N*_*beads*_, to generate loops of realistic dimensions. Similarly, the coefficient *b* must be selected to align with this order of magnitude to effectively balance the impact of the entropic cost in the energy. Moreover, *N*_(*·*)_ represents the normalization constants for each factor,

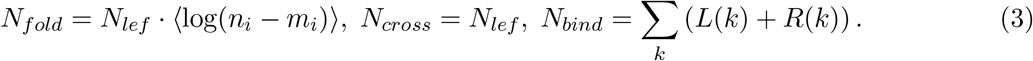

Therefore, the energy difference can be written as a sum of three factors,

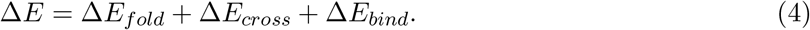

In this manner we accept a move in two cases:

1. If Δ*E <* 0 or,
2. if Δ*E >* 0 with probability *e*^*−*Δ*E/kT*^.

The resulting equilibrium distribution of loops depends on the temperature *T* of the Boltzmann distribution. This process represents the Metropolis stochastic simulation of LoopSage. Alternatively, the Simulated Annealing method provides an additional option for modeling, where the temperature decreases according to *T*_*k*_ = (1 − *k/N*_steps_) *T*_0_.

After running a stochastic simulation, we obtain a set of LEF positions (*m*_*i*_(*t*), *n*_*i*_(*t*)) as the outcome (see Fig.2a). The main advantage of this simulation is that LEFs binding at convergent CTCF sites tend to be more stable and have longer lifetimes, as the system reaches a local minimum. This behavior aligns with experimental knowledge indicating that LEFs exhibit increased stability when encountering a BE [42, 43]. We determine the simulation equilibrium by monitoring the system’s energy (see Fig.2b). The simulation is considered to be in equilibrium when the energy maintains an approximately constant value after the burn-in period. Additionally, we define a sampling step to ensure that ensembles of LEF positions are independently and identically distributed (see Fig.2c).

**Figure 2:**
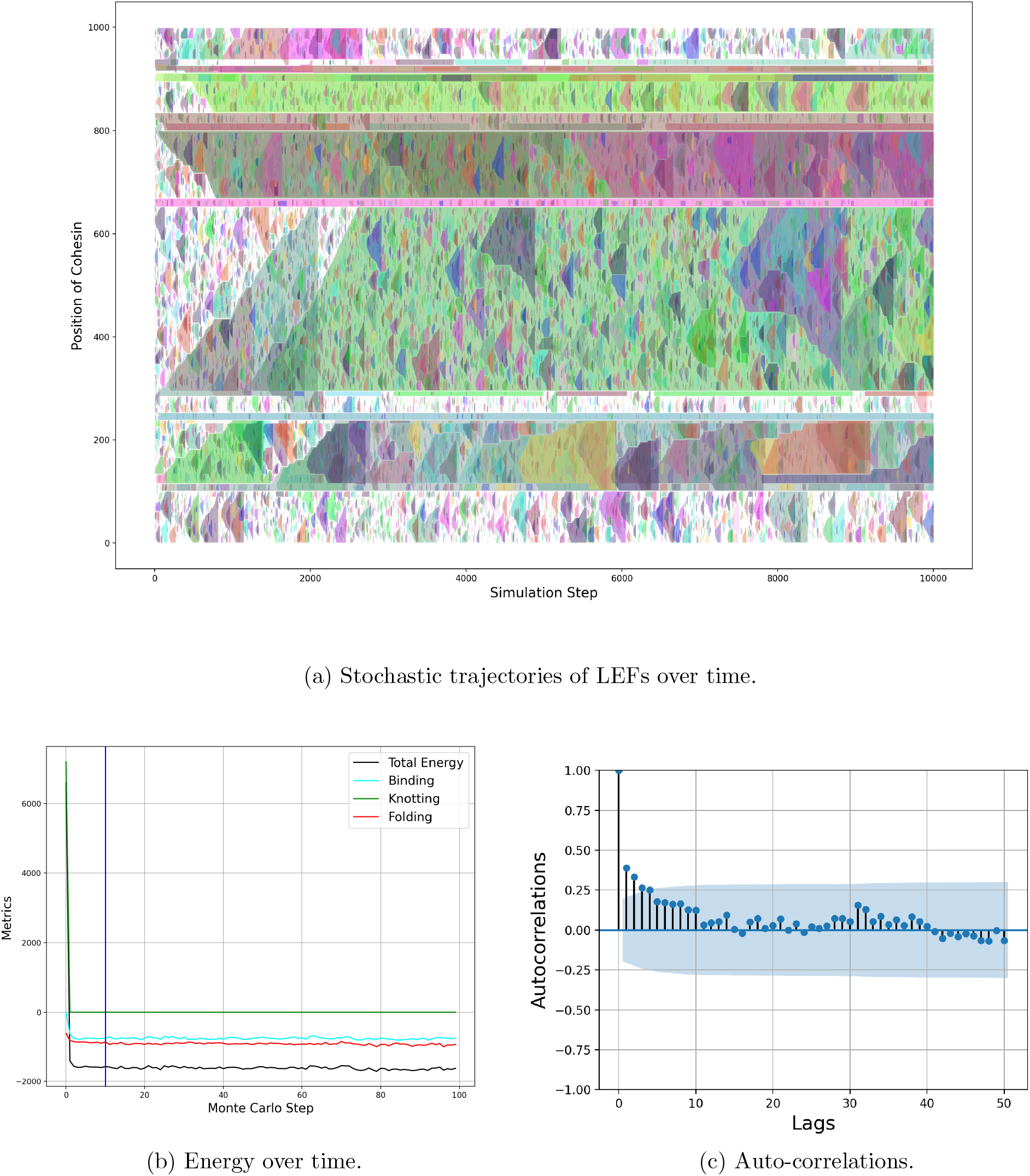
An example run of stochastic simulation LoopSage for loops from CTCF ChIA-PET data, for the region [225236830, 226046745] of chromosome 1 for the celline GM12878. In this example we see a simulated annealing stochastic simulation with initial temperature *T* = 5, Monte Carlo steps *N*_*steps*_ = 10^4^, Monte Carlo step *s* = 10^2^ and burnin period *bi* = 1000. In this simulation the number of beads is *N*_*beads*_ = 1000, the number of LEFs *N*_*lef*_ = 50, and the coefficients of the energy *f* = *−*1000, *b* = *−*1000 and *κ* = 20000. (a) The trajectories of LEFs as a function of time. The extruded area of each LEF is represented with different color. In y-axis there is the linear polymer distance, whereas in x-axis there is Monte Carlo time. (b) The evolution of energy as a function of time. With blue vertical line burn-in period is represented. (c) Auto-correlations.

### 2.3 Molecular Dynamics Simulation

Let us consider a system comprising *N*_lef_ LEFs, as well as two matrices, *M* and *N*, both of which possess dimensions *N*_coh_ × *N*_steps_. These matrices represent the respective constraints associated with each LEF. Consequently, we define a time-dependent force field as follows:

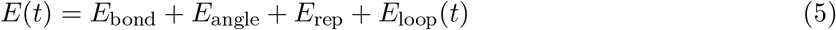

where,

- *E*_bond_ corresponds to a typical harmonic bond force field that connects adjacent beads *i* and *i* + 1, with an equilibrium length of *x*_0_ = 0.1 nm and a Hook constant assumed to be *k* = 3 × 10^5^ kJ/(mol · nm^2^).
- *E*_angle_ a harmonic angle force that connects beads *i* − 1, *i, i* + 1, and has equilibrium angle *θ*_0_ = *π* and Hook constant 400 *kJ/*(*mol* · *nm*^2^).
- *E*_rep_ which is a repelling forcefield of the form:

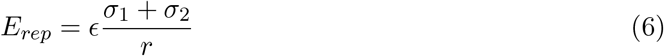

where *ϵ* = 0.5 *kJ/mol* and *σ*_1_ = *σ*_2_ = 0.05 *nm*.
- *E*_loop_(*t*) represents a time-dependent loop extrusion force. This force reads the matrices *M* and *N*, applying a distinct set of constraints C_*ti*_ = {*m*_*j*_(*t*_*i*_), *n*_*j*_(*t*_*i*_)} at each time step *t*_*i*_. Each LEF *j* is subjected to specific constraints *m*_*j,ti*_ and *n*_*j,ti*_ . The functional form of this force is also a harmonic bond force, with parameters *x*_0_ = 0.1 nm and a Hook constant assumed to be *k* = 3 × 10^4^ kJ/(mol · nm^2^).

For the implementation of this model in python, we used OpenMM [44, 45] and CUDA acceleration. To minimize the energy Langevin dynamics [46] were used, in temperature of *T*_*L*_ = 310 *K*, friction coefficient *γ* = 0.05 *psec*^*−*1^ and time step *t*_*s*_ = 100 *fsec*. Note that the temperature of molecular dynamics simulation is independent from the temperature of stochastic simulation and they represent different physical realities.

### 2.4 Connecting the dots

As previously discussed, the primary objective of this simulation pipeline is to reconstruct experimental ChIA-PET heatmaps by exploring statistical ensembles of 3D structures. To achieve this, a combined approach involving both stochastic and molecular dynamics simulations is employed (Fig.1D). Initially, a minimal set of data is required, including positions of Bes and orientations, which are used to determine the energy minima of the simulations. The energy function *L*(*m*_*i*_) is defined as follows:

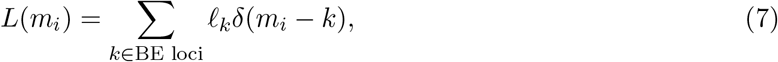

where *ℓ*_*k*_ = _*i*_ *c*_*ki*_ · *p*_left,*ki*_. Here, *c*_*ki*_ represents the PET-count between loop *k* and *i*, and *p*_left,*ki*_ denotes the probability of the CTCF motif having a tandem-left orientation. Similarly, *R*(*n*_*i*_) is defined. Typically, BE positions can be determined using ChIA-PET loop anchors in the .bedpe format, with loops containing a PET-count greater than 10 being retained. Optionally, a ChIP-Seq file can be imported if the user desires preferential loading of LEF in enriched regions.

The imported data is then used in the stochastic simulation pipeline to generate the stochastic trajectories of LEFs (*m*_*i*_(*t*), *n*_*i*_(*t*)) (Fig.1D(i)). These trajectories are subsequently incorporated into the molecular dynamics pipeline, implemented using OpenMM, to generate independent and identically distributed (iid) ensembles of structures (Fig.1D(ii)). From each of these ensembles, the all-versus-all inverse distance heatmap is computed, resulting in an ensemble of heatmaps (Fig.1D(iii)). Finally, the experimental ChIA-PET heatmap is reconstructed by averaging all the simulation heatmaps (Fig.1D(iv)).

## 3. Results

### 3.1 Parameter Study

Firstly, we examine the bifurcations within the stochastic simulation pipeline. It is noteworthy that, according to our modeling method, stochastic simulation can operate independently of molecular dynamics, thereby representing distinct physical realities. The physical systems described by stochastic simulation encompass ensembles of loop conformations, corresponding to LEF positions. The thermodynamic properties of the final conformation ensemble are heavily reliant on simulation parameters. Hence, conducting a parameter study prior to modeling real regions is crucial. In this section, we focus on two statistical metrics that characterize the macrostates of the system. The one it is the folding of chromatin as it is already defined by the first term of eq.1, and the other one is the proportion of gaps in the polymer, defined as follows,

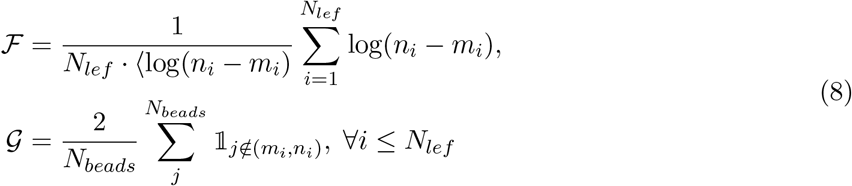

It is notable that these metrics are normalized in such a way that the final result does not depend a lot on the polymer length *N*_*beads*_ and this makes the parameter analysis easier, because there is one less parameter to study.

Within this model, we consider five parameters of interest: the folding coefficient *f*, the crossing coefficient *k*, the binding coefficient *b*, the temperature *T*, and the ratio of LEFs to BEs *N*_*lef*_ */N*_*be*_. However, we can effectively reduce these parameters to four by assuming an infinite crossing coefficient *k* = ∞. Although understanding the phase transitions of a system with numerous parameters is challenging, it is still feasible to generate diagrams that depict how the average folding of chromatin depends on system parameters.

The first observation from plotting the folding and the proportion of gaps as functions of temperature (Fig. 3) is the existence of a phase transition in the system between a condensed and a sparse state. This transition is determined by the system’s temperature *T* and the number of LEFs in the simulation. When the LEF count is small, a peak in the lower temperature range is observed, as LEFs have sufficient time to form larger loops and abundant free space to occupy. Conversely, when the LEF count is higher, the proportion of gaps decreases. This is attributed to the system being LEF-rich, resulting in smaller loops being present throughout the region because LEFs block each other due to the crossing penalty (Fig.1B). For simulations with a lower LEF count, a phase transition in the proportion of gaps occurs. Specifically, at low temperatures, the condensed state is characterized by a minimal number of gaps, while at higher temperatures, LEFs rapidly relocate, leading to the formation of predominantly small loops and a higher proportion of gaps, indicating the sparse state. Furthermore, the values of the parameters *f* and *b*, representing the folding and binding energies, respectively, play a significant role in loop extrusion dynamics. For instance, in Fig. 3a, the temperature interval associated with higher folding occurs between temperatures *T* ∈ [0, 10]. Increasing the values of *f* and *b* from *f* = *b* = −500 expands this temperature interval, as higher folding and binding coefficients induce faster condensation of the system (Fig. 3d).

**Figure 3:**
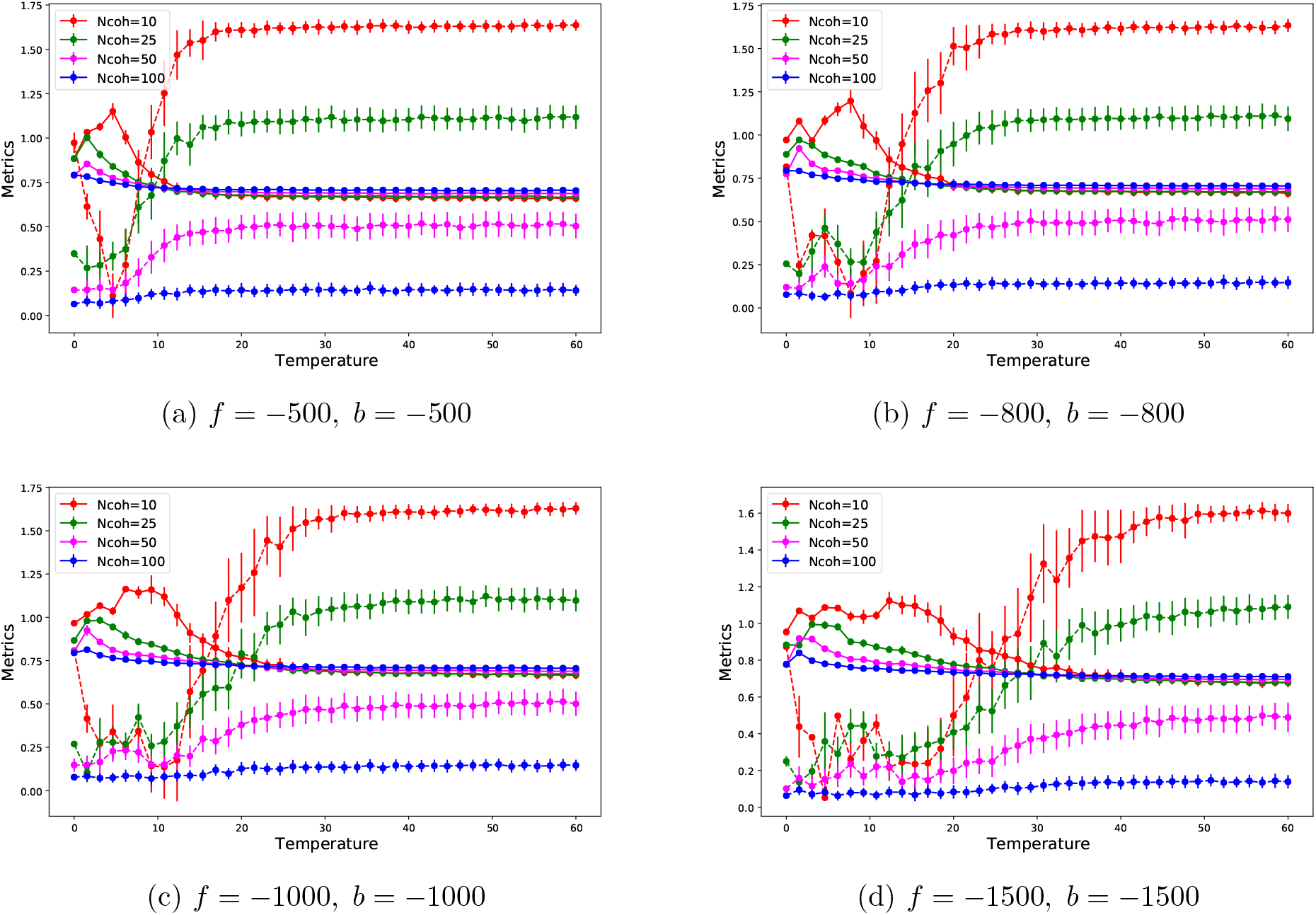
Diagram of folding (continuous line) and the proportion of gaps (dashed line) under the change of temperature. In these simulations we set the parameters *N*_*beads*_ = 1000, *κ* = 100000. We iterate over 2000 Monte Carlo steps with sampling frequency *t*_*s*_ = 100 and burnin period *bi* = 1000. We simulated 40 different values of temperature *T* in the interval [0, 60] for each number of LEFs. The number of BEs in the system is *N*_*be*_ = 44. The modelling region that we used as reference was the region [178421513, 179491193] of chromosome 1.

Finally, it would be interesting to check the behavior of our system when *κ* coefficient changes. Interestingly, it is observed that for lower values of *κ*, LEFs are capable of passing through each other and this may lead to bigger loop sizes. Therefore, Fig. 4 shows that for lower values of *κ*, folding metric has slightly higher values. As it is expected by intuition, the proportion of LEFs in penalized configurations of Fig.1B is increased as the number of LEFs is increased.

**Figure 4:**
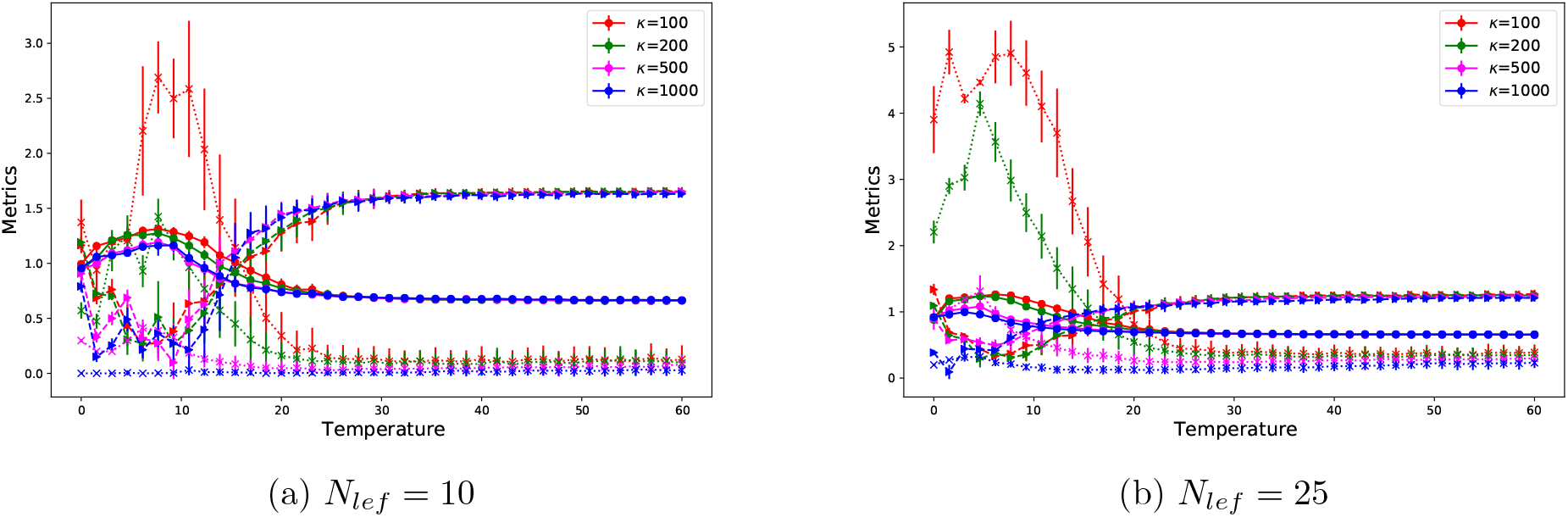
Diagram of folding (continuous line) and the proportion of gaps (dashed line) under the change of temperature. In these simulations we set the parameters *N*_*beads*_ = 1000, *f* = *b* = *−*1000. We iterate over 10000 Monte Carlo steps with sampling frequency *t*_*s*_ = 100 and burnin period *bi* = 1000. We simulated 40 different values of temperature *T* in the interval [0, 60] for each *κ*. The number of BEs in the system is *N*_*be*_ = 44. The modelling region that we used as reference was the region [178421513, 179491193] of chromosome 1. With marker ‘o’ we represent the folding metric of chromatin, with the marker ‘*>*’ we represent the proportion of gaps and with ‘*×*’ we represent the proportion of LEFs with penalized configurations.

### 3.2 Modelling experimental data

LoopSage demonstrated remarkable efficiency in modeling experimental heatmaps. To simulate TAD regions, our approach requires a minimal set of data: (i) BEs positions and orientations, and (ii) cohesin coverage tracks. For BE positioning, we utilize ChIA-PET interaction data, which we filter to determine the probabilities of forward or backward orientations. Cohesin data provide information on LEF enrichment in different loci within the modeling region. Interestingly, we discovered that the precise information regarding LEF enrichment is not critical for reconstructing 3C heatmaps. This is because LEFs, owing to their biophysical properties, exhibit greater enrichment in CTCF-enriched regions. Consequently, based on our modeling, LEFs naturally spend more time in BE energy gaps. Thus, we can choose to either preferentially load LEFs in regions with observed enrichment in ChIPSeq data or explore the phase space more homogeneously. However, such choices do not significantly alter the resulting heatmaps. Accordingly, we did not use ChIP-Seq data for the presentation of results of this study.

Our research highlights an alternative, faster approach to reconstructing heatmaps, distinct from the conventional method of calculating all-versus-all distances of 3D structures (Fig.5c). We initially plot all LEF positions obtained from the stochastic simulation output for a given time point. Next, we create a set of artificial loops based on the communication property of single molecule structures. According to this property, if a loop connects locus *a* with locus *b*, and another loop connects *b* with *c*, there must also be a loop between *a* and *c* due to the geometry of the problem. We plot these artificial loops on our heatmap and fill in the squares between the loops. The intensity on the heatmap corresponds to the number of loops present in a particular region. From this perspective, 3C experiment heatmaps are composed of two distinct sets of loops: short-range loops resulting from LEF bonds, and long-range loops arising from the communication property between adjacent short-range loops. This approach enables the rapid reproduction of experimental heatmaps without the need for molecular simulations. However, it is important to note that certain information associated with the 3D geometry of chromatin may be lost in this process.

**Figure 5:**
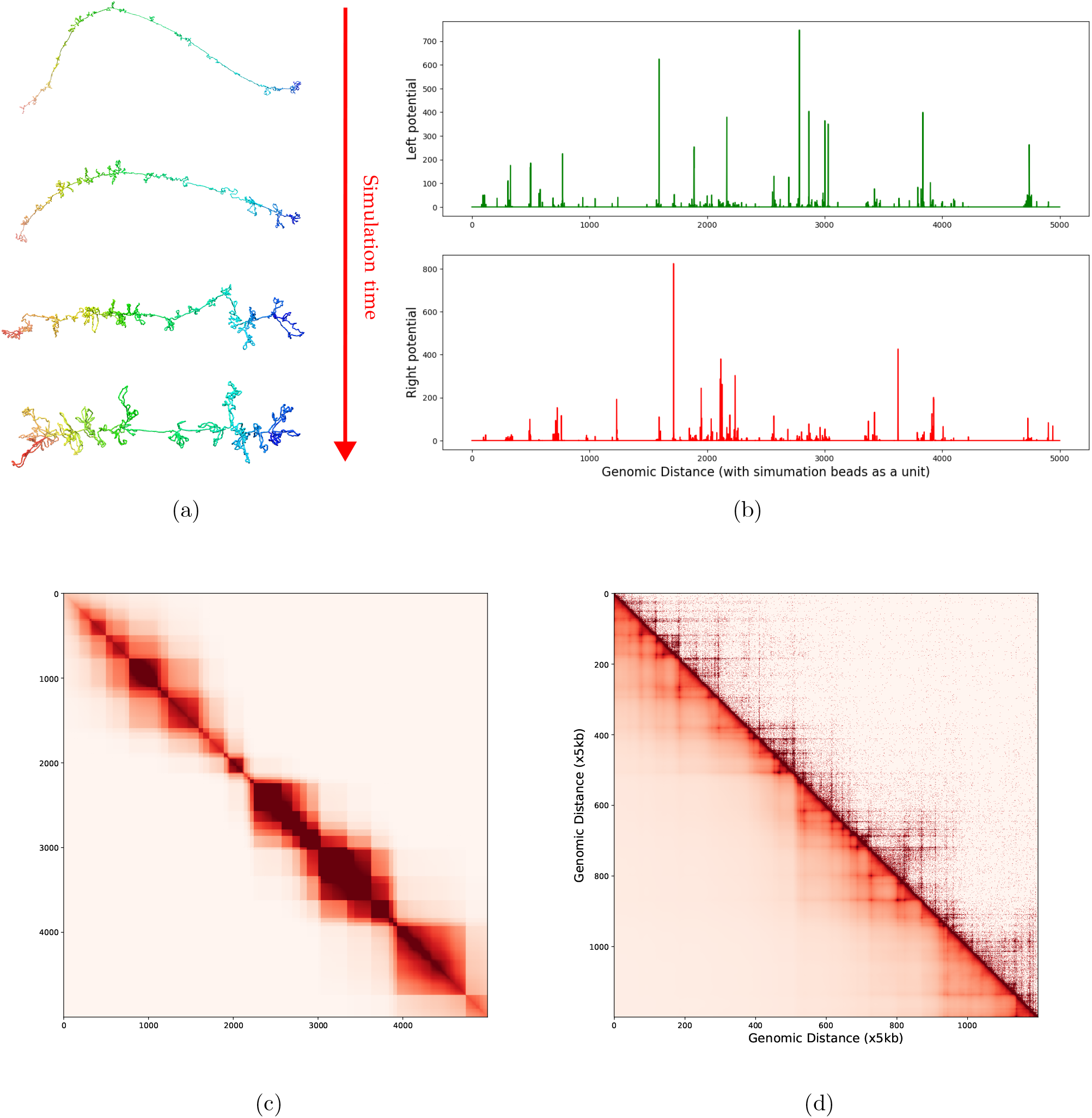
Simulating region [165000000, 171000000] of chromosome 1. For the purposes of this simulation we used *N*_*beads*_ = 5000, with energy parameters *f* = *−*6000, *b* = *−*3000 and *k* = 10^5^. For the stochastic simulated annealing simulation *N*_steps_ = 5000 were performed, with frequency *t*_*s*_ = 100 and burn-in period *bi* = 1000. We can see the following figures: (a) How the 3D-structure changes as a function of simulation time. Initially, small loops are formed in the arbitrary chosen initial structure and it becomes more and more compacted with bigger loops being formed. (b) Left and right binding potential that it is used as input and it is computed by the CTCF motif strength and probability to be in forward or backward orientation. (c) Heatmap that it is computed only from LEF positions after stochastic simulation without running dynamical molecular simulation. (d) Comparison between experimental CTCF ChIA-PET and simulation heatmap for the given region. The Pearson correlation between the two heatmaps is 0.746.

To address this limitation, our pipeline incorporates the option of molecular simulations. LEF positions are modeled as time-dependent covalent harmonic bonds, and our OpenMM script calculates the stochastic 3D trajectory of the chromatin region Fig.5a. Ensembles of structures are generated by saving structures at specific Monte Carlo steps. Initially, small loops are formed within an arbitrary initial structure, typically a self-avoiding random walk. After a burn-in period, a statistical ensemble of single molecule structures can be obtained. These structures exhibit complex shapes composed of both larger and smaller loops. In the case of simulated annealing, we consider the final structure as the most representative, while in the case of Metropolis simulation, a statistical analysis is performed to identify the most representative structure. A quaternion RMSD heatmap [47–49] is constructed to measure the similarities between different structures. By applying multidimensional scaling (MDS) dimensionality reduction [50] to this heatmap, followed by DBSCAN clustering [51, 52], we identify the structure within the ensemble that is closest to the centromere as the most representative structure.

The validation of our results proved challenging due to the inherent differences between experimental and simulation heatmaps. However, as shown in Fig. 6, which presents the modeling of four distinct TAD regions from chromosomes 1 and 2, we observe significant similarities in stripe and loop patterns between the simulation and experimental data. To further evaluate these patterns of interest (i.e., loops and stripes), we propose a metric based on comparing ChIA-PET interactions and loop patterns in the simulation heatmaps. Specifically, we aggregate the PET count exclusively in loci with CTCF loops by summing in the second dimension of the experimental heatmap. The same procedure is applied to the simulation heatmaps, resulting in two signals that can be readily compared using statistical measures such as Pearson correlation. Employing this approach, we find substantial similarities (over 75%) between the resulting heatmaps and experimental data.

**Figure 6:**
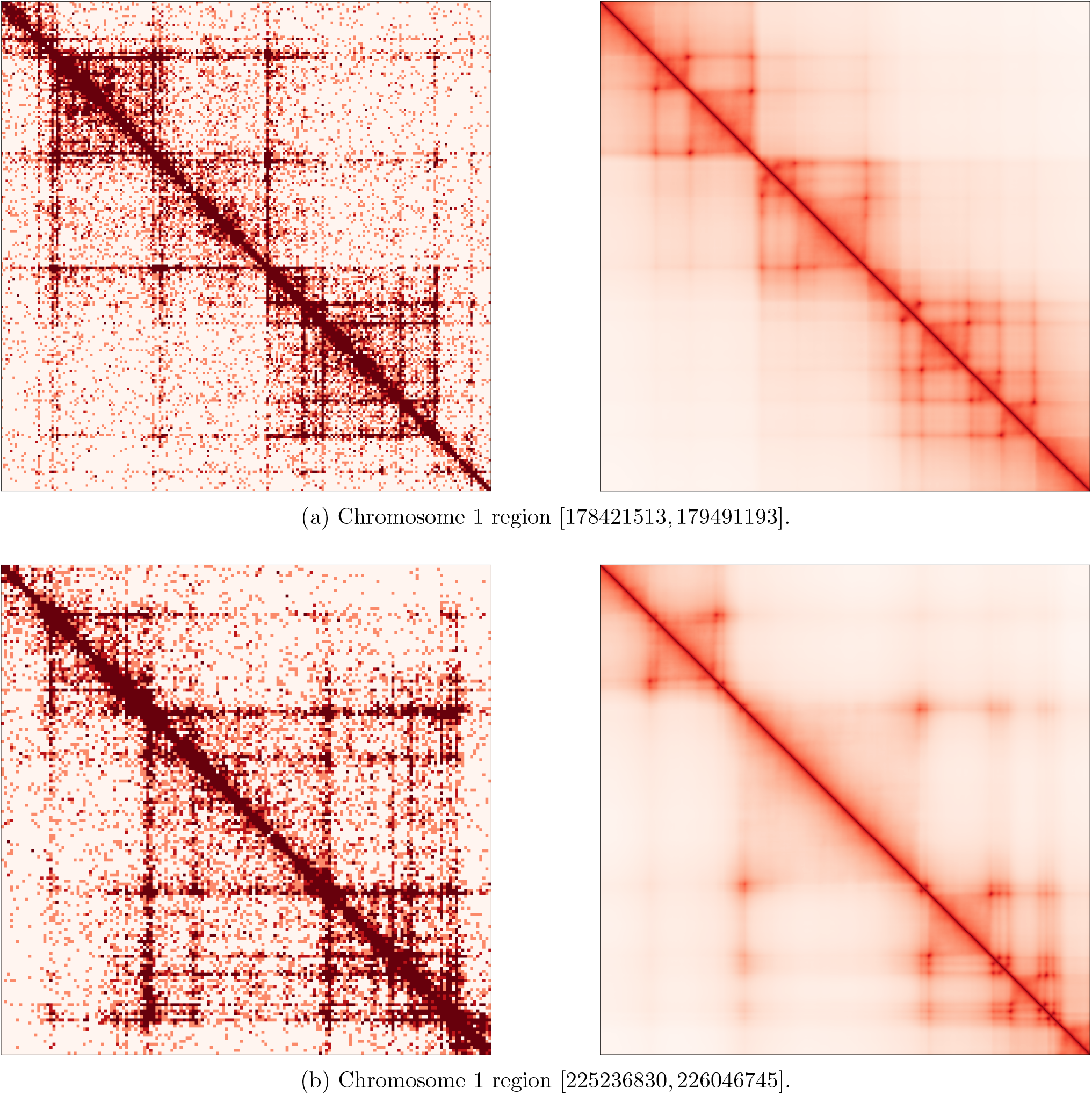
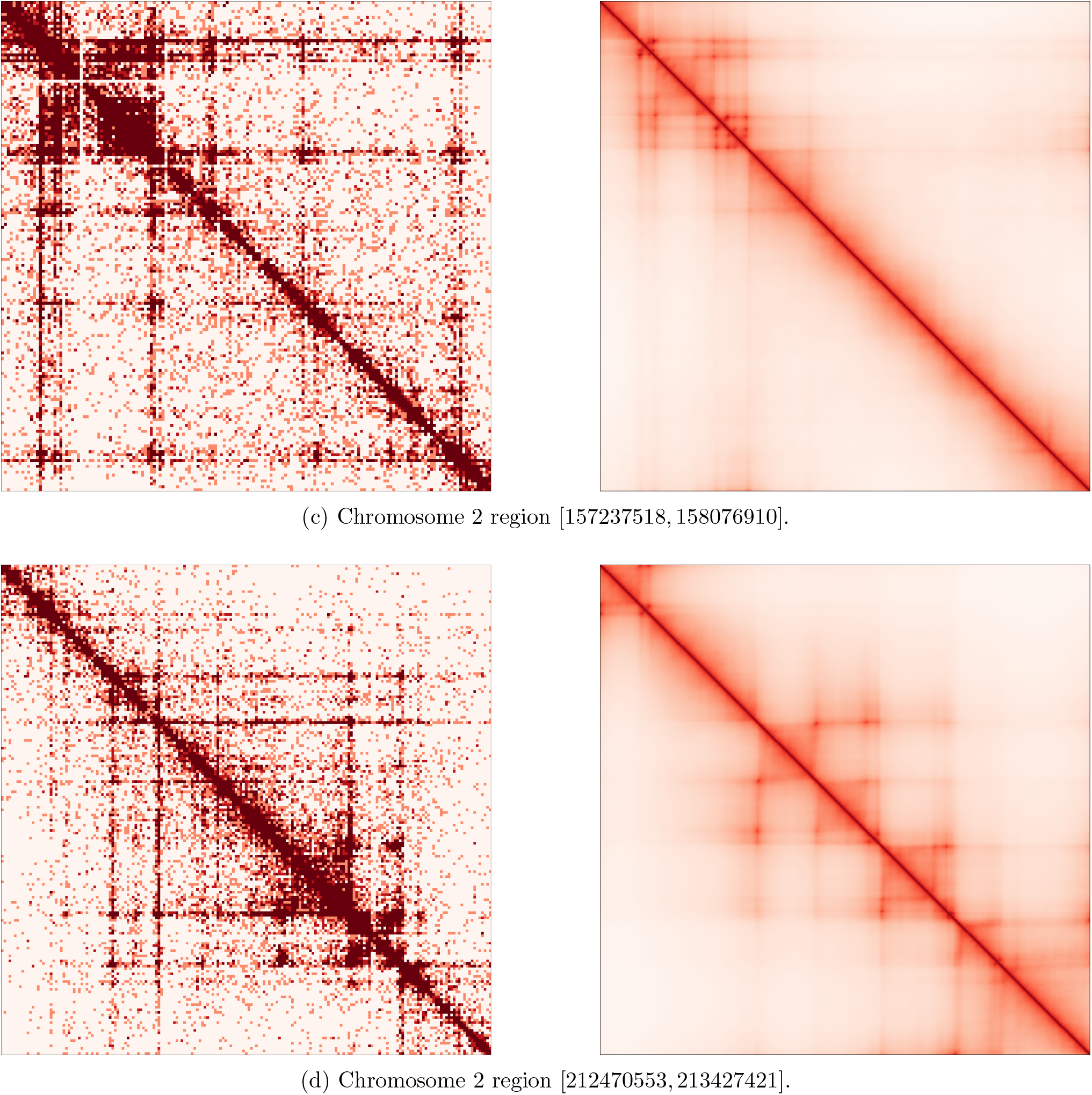
On the left-hand-side CTCF ChIA-PET heatmaps, and on the right-hand-side result heatmaps of stochastic simulation LoopSage for loops from CTCF ChIA-PET data, for the given regions of celline GM12878. In this example we see a simulated annealing stochastic simulation with initial temperature *T* = 5, Monte Carlo steps *N*_*steps*_ = 10^4^, Monte Carlo step *s* = 10^2^ and burnin period *bi* = 1000. In this simulation the number of beads is *N*_*beads*_ = 1000, the number of LEFs *N*_*lef*_ = 50, and the coefficients of the energy *f* = *−*1000, *b* = *−*1000 and *κ* = 20000. The following Pearson correlations between simulated and experimental heatmaps are estimated: (a) 78%, (b) 91.6%, (c) 76.1%, and (d) 81.1% .

## 4. Discussion

In this study, we have implemented an innovative approach for loop extrusion modeling, which effectively reconstructs experimental ChIA-PET heatmaps. While there exist similar works [13, 15, 17, 19] that utilize stochastic simulations and Monte Carlo methods, our approach stands out as it is the first to leverage a Hamiltonian function constructed based on first-principle biophysical assumptions associated with LEFs motion and its interaction with BEs. The mathematical elegance of our approach allows us to provide readers with a simplified Python script capable of generating realistic single-molecule structures by translating LEF positions into covalent bonds. This enables comparisons between theoretical models and experimental data. Moreover, our stochastic 3D trajectory computation is highly efficient, typically taking only a few minutes, thanks to the acceleration provided by CUDA graphics card computing within the OpenMM software framework [44, 45]. It is worth noting that we achieved our goal using a minimal set of data primarily consisting of CTCF anchors. However, our modeling framework can accommodate a generic input approach, allowing users to construct binding potentials as desired.

Another important advancement, in contrast to previous studies, is the improved formulation of phase transitions within our model. Consequently, the occurrence of dense and sparse chromatin states emerges naturally as a result of changes in Hamiltonian parameters and temperature within the Boltzmann distribution. This new perspective adds depth to prior investigations that sought to model loop extrusion as a physical system with phase transitions [32–36]. The main advantage of this method is that the stochastic simulation is performed in the one dimensional DNA fiber, and it is independent of molecular dynamics. This approach illustrates a new connection between different dimensionalities of chromatin folding and it connects two different approaches of modelling: stochastic simulation and energy based modelling with phase transitions. From a biological standpoint, our model incorporates current knowledge from live-imaging studies regarding the stability of CTCF loops [43], successfully reproducing single-molecule structures at the scale of topologically associated domains (TADs). In summary, our method serves a dual purpose: assisting biologists in understanding chromatin structure and providing a theoretical framework for loop extrusion that may inspire physicists to explore phase transitions in chromatin.

Despite the success of our method, there remain open questions concerning the stochastic modeling of loop extrusion. One important aspect is the tuning of hyperparameters in the simulation to produce the best possible results, ensuring that the generated heatmaps closely resemble the experimental ones. Further improvements in this type of modeling could involve the inclusion of other obstacles, such as nucleosome aggregates, RNA polymerase II, MCM complexes, as proposed in previous studies [21–23].

To conclude, our study shows a way to simulate stochastically LEF trajectories by making hypotheses that are linked to the single cell biology. Therefore, it brings a new light to the previous knowledge that we had about the loop, stripe and TAD patterns of 3C experiments, since instead of thinking them as family of loops averaged among the population of cells, we see that these patterns occur naturally from the stochastic dynamics of loop extrusion. The formalism of our method gives an elegant and simple way to simulate such regions, and it adds a refreshed look in the modelling of LEF dynamics connecting the one dimensional DNA fiber with the three dimensional folded chromatin structure.

## Data availability

For this project publicly available data from ENCODE were used. The discussed results are based on the data of Yijun Ruan, JAX laboratory for CTCF-ChIA-PET of the celline GM12878 (project ENCSR184YZV).

## Code Availability

The code of LoopSage is available as an open-source project on the GitHub repository https://github.com/SFGLab/LoopSage. Furthermore, CTCF Motif search script is available on the GitHub as an open-source project as well https://github.com/SFGLab/3d-analysis-toolkit. Please, cite this article in any work that uses our code.

## Acknowledgements

Research was funded by Warsaw University of Technology within the Excellence Initiative: Research University (IDUB) programme. This work has been co-supported by Polish National Science Centre (2019/35/O/ST6/02484 and 2020/37/B/NZ2/03757). Computations were performed thanks to the Laboratory of Bioinformatics and Computational Genomics, Faculty of Mathematics and Information Science, Warsaw University of Technology using Artificial Intelligence HPC platform financed by Polish Ministry of Science and Higher Education (decision no. 7054/IA/SP/2020 of 2020-08-28). We would like to thank our laboratory member Mateusz Chiliński who helped us in the CTCF motif search task.

## Notes

https://github.com/SFGLab/LoopSage

### Competing Interest Statement

The authors have declared no competing interest.

